## Supplementary figures and small tables (PDF) (v3) for "Unexpected mutual regulation underlies paralogue functional diversification and promotes maturation of a protective epithelial tissue"

### Supplementary Material

Within this PDF:

Figure S1. Alignments of *zen1* and *zen2* homeoboxes/domains.

Figure S2. Additional phylogenies of insect class 3 Hox proteins.

Figure S3. Phenotypic consequences of *Tc-zen2* RNAi - milder defects.

Figure S4. Tc-Zen1 and Tc-Zen2 expression detected by western blotting.

Figure S5. Early *in vivo* expression dynamics of transcript and protein for the *Tc-zen* paralogues.

Figure S6. Expression screen summary of early Tc-Zen2 candidate targets.

Figure S7. Gene ontology (GO) enrichment analysis after *Tc-zen2* RNAi.

Figure S8. Functional annotation and validation of late Tc-Zen2 targets.

Table S5. List of gene ontology (GO) terms grouped by category of interest in each GO domain.

Table S6. Description of Tc-Zen2 late candidate target genes validated by RT-qPCR.

Table S7. *Tribolium castaneum* (*TC*) primer sequences for *in situ* hybridization, RNAi, and RT-qPCR.

Table S8. Comparison of alignment statistics by read length.

Excel workbook file (large datasets and gene lists):

| Table | Description |
|---|---|
| S1A | List of differentially expressed genes during early development after *Tc-zen1* RNAi ($P_{adj}$≤0.01) |
| S1B | List of differentially expressed genes during early development after *Tc-zen2* RNAi ($P_{adj}$≤0.01) |
| S1C | List of differentially expressed genes during early development (WT *Tc-zen1* peak, 6-10 hAEL vs. WT *Tc-zen2* peak, 10-14 hAEL) ($P_{adj}$≤0.01) |
| S2A | Regulatory targets shared by both *Tc-zen* paralogues with relaxed thresholds ($P_{adj}$≤0.05): same direction of regulation, N=42 |
| S2B | Regulatory targets shared by both *Tc-zen* paralogues with relaxed thresholds ($P_{adj}$≤0.05): opposite direction of regulation, N=78 |
| S3A | List of differentially expressed genes during late development (pre-rupture, 48-52 hAEL) after *Tc-zen2* RNAi ($P_{adj}$≤0.01) |
| S3B | List of differentially expressed genes during late development (rupture/post-rupture, 52-56 hAEL) after *Tc-zen2* RNAi ($P_{adj}$≤0.01) |
| S3C | List of differentially expressed genes during late development (WT pre-rupture, 48-52 hAEL vs. WT rupture/post-rupture, 52-56 hAEL) ($P_{adj}$≤0.01) |
| S3D | List of differentially expressed genes during late development (*Tc-zen2* RNAi pre-rupture, 48-52 hAEL vs. *Tc-zen2* RNAi rupture/post-rupture, 52-56 hAEL) ($P_{adj}$≤0.01) |
| S3E | List of differentially expressed genes during late development (WT pre-rupture, 48-52 hAEL vs. *Tc-zen2* RNAi rupture/post-rupture, 52-56 hAEL) ($P_{adj}$≤0.01) |
| S4A | Enriched GO terms in the dataset "Differentially expressed genes during pre-rupture (48-52 hAEL stage) after *Tc-zen2* RNAi" |
| S4B | Enriched GO terms in the dataset "Differentially expressed genes during rupture/post-rupture (52-56 hAEL stage) after *Tc-zen2* RNAi" |

A

```
        1                                   10                                      20
                                                         HELIX 1
        G   K   R   A   R   T   A   Y   T   S   A   Q   L   V   E   L   E   R   E   F
        G   K   R   A   R   T   A   Y   T   S   A   Q   L   V   E   L   E   R   E   F
        G   K   R   A   R   T   A   Y   T   S   A   Q   L   V   E   L   E   R   E   F
        P   K   R   V   R   T   A   Y   T   S   A   Q   L   V   E   L   E   R   E   F
Tcas-z1 GGA AAG CGT GCC CGC ACT GCC TAC ACC TCG GCC CAA TTA GTG GAA TTA GAG CGG GAA TTC
Tfre-z1 GGA AAA CGT GCT CGC ACT GCT TAC ACC TCA GCC CAG TTA GTG GAA TTA GAG CGG GAA TTC
Tmad-z1 GGA AAA CGT GCT CGC ACT GCG TAT ACC TCG GCT CAA TTA GTC GAA TTA GAG CGG GAA TTT
Tcon-z1 CCA AAA CGT GTC CGC ACT GCT TAC ACA TCA GCC CAA TTA GTA GAA CTA GAA CGG GAA TTC
         *  **  *** *   *** *** **  **  **  **  **  **  *** **  ***  ** **  *** *** **

        21                                  30                                      40
                                                 HELIX 2
        H   H   G   K   Y   L   S   R   P   R   R   I   Q   I   A   E   N   L   N   L
        H   L   G   K   Y   L   S   R   P   R   R   I   Q   M   A   E   N   L   N   L
        H   R   G   K   Y   L   T   R   P   R   R   I   Q   M   A   E   N   L   K   L
        H   R   G   K   Y   L   S   R   P   R   R   I   Q   M   A   T   N   L   N   L
Tcas-z1 CAC CAC GGG AAA TAC TTA TCA CGA CCA CGC CGA ATC CAA ATC GCC GAG AAC CTG AAC CTG
Tfre-z1 CAT CTC GGG AAA TAT TTA TCA CGG CCG CGA CGA ATC CAA ATG GCC GAA AAT CTA AAC CTG
Tmad-z1 CAT CGC GGG AAA TAT TTA ACA CGG CCG CGA AGA ATC CAA ATG GCT GAA AAT TTG AAA CTG
Tcon-z1 CAT CGT GGA AAA TAT TTA TCA CGA CCT AGG AGA ATC CAG ATG GCC ACA AAT CTA AAT CTC
        **  *   **  *** **  ***  ** **  **   *   ** *** **  **  **      **   *  **  **

        41                                  50                                      60
                                     HELIX 3
        S   E   R   Q   I   K   I   W   F   Q   N   R   R   M   K   H   K   K   E   Q
        S   E   R   Q   I   K   I   W   F   Q   N   R   R   M   K   Y   K   K   E   Q
        S   E   R   Q   I   K   I   W   F   Q   N   R   R   M   K   H   K   K   E   Q
        S   E   R   Q   I   K   I   W   F   Q   N   R   R   M   K   H   K   K   E   Q
Tcas-z1 AGT GAG CGT CAG ATC AAG ATT TGG TTC CAA AAT CGC AGA ATG AAG CAC AAG AAA GAA CAG
Tfre-z1 AGT GAG CGT CAA ATC AAA ATT TGG TTC CAG AAC CGA AGA ATG AAG TAC AAG AAG GAG CAG
Tmad-z1 AGT GAA CGT CAA ATC AAA ATT TGG TTC CAA AAT CGA AGA ATG AAG CAC AAG AAA GAA CAG
Tcon-z1 AGT GAA CGT CAA ATC AAG ATT TGG TTT CAA AAT CGC AGA ATG AAG CAC AAG AAG GAG CAA
        *** **  *** **  *** **  *** *** **  **  **  **  *** *** ***  ** *** **  **  **
```

B

```
        1                                   10                                      20
                                                         HELIX 1
        G   K   R   A   R   T   A   Y   T   S   S   Q   L   V   E   L   E   R   E   F
Tcas-z2 GGC AAA AGG GCC CGA ACT GCG TAC ACC TCC TCC CAA TTA GTC GAA CTC GAA CGC GAA TTT
Tfre-z2 GGC AAA AGG GCC CGA ACT GCA TAC ACC TCC TCT CAA TTA GTC GAA CTG GAA CGC GAA TTT
Tmad-z2 GGC AAA AGA GCC CGA ACT GCC TAC ACC TCC TCA CAA TTA GTA GAA TTA GAA CGC GAA TTT
Tcon-z2 GGA AAA AGA GCT CGA ACT GCA TAC ACT TCC TCT CAA TTA GTC GAA TTA GAG CGC GAA TTC
        **  *** **  **  *** *** **  *** **  *** **  *** *** **  ***  *  **  *** *** **

        21                                  30                                      40
                                                 HELIX 2
        H   R   S   K   Y   L   C   R   P   R   R   I   Q   M   A   Q   N   L   N/S L
Tcas-z2 CAC CGG AGC AAG TAC TTG TGC CGC CCT CGG CGC ATC CAA ATG GCC CAA AAC CTC AAC TTG
Tfre-z2 CAC CGC AGT AAA TAC TTA TGT CGA CCT CGG CGC ATM CAA ATG GCC CAA AAT CTC AAC TTG
Tmad-z2 CAT CGA AGC AAA TAT TTA TGT CGT CCT CGT CGT ATT CAA ATG GCC CAA AAC CTC AAT TTA
Tcon-z2 CAC CGT AGC AAG TAT TTA TGT AGA CCT CGT CGC ATC CAA ATG GCT CAA AAT CTA AGC CTT
        **  **  **  **  **  **  **   *  *** **  **  **  *** *** **  *** **  **  *    *

        41                                  50                                      60
                                     HELIX 3
        T   E   R   Q   I   K   I   W   F   Q   N   R   R   M   K   F   K   K   E   E
Tcas-z2 ACC GAA CGC CAA ATC AAG ATT TGG TTC CAA AAC CGC AGA ATG AAG TTC AAA AAG GAG GAG
Tfre-z2 ACT GAG CGT CAA ATC AAG ATT TGG TTC CAA AAC CGT AGA ATG AAG TTC AAA AAG GAG GAA
Tmad-z2 ACT GAA CGC CAA ATA AAA ATT TGG TTT CAA AAC CGA CGT ATG AAG TTC AAG AAA GAA GAA
Tcon-z2 ACT GAA CGC CAA ATC AAA ATT TGG TTT CAA AAT CGT CGC ATG AAA TTC AAA AAA GAA GAA
        **  **  **  *** **  **  *** *** **  *** **  **   *  *** **  *** **  **  **  **
```

codons with non-synonymous substitutions    M A or C (nucleotide sequence ambiguity)

Nucleotide substitution at: first position second position third position

**Figure S1. Alignments of *zen1* and *zen2* homeoboxes/domains.** The alignments of *zen1* **(A)** and *zen2* **(B)** homeoboxes/domains of four *Tribolium* species (*T. castaneum* - *Tcas*; *T. freeman* - *Tfre*; *T. madens* - *Tmad*; *T. confusum* - *Tcon*) show that the number of non-synonymous substitutions is higher in the Zen1 homeodomain (9 substitutions) than in the Zen2 homeodomain (1; see also Fig. 1B).

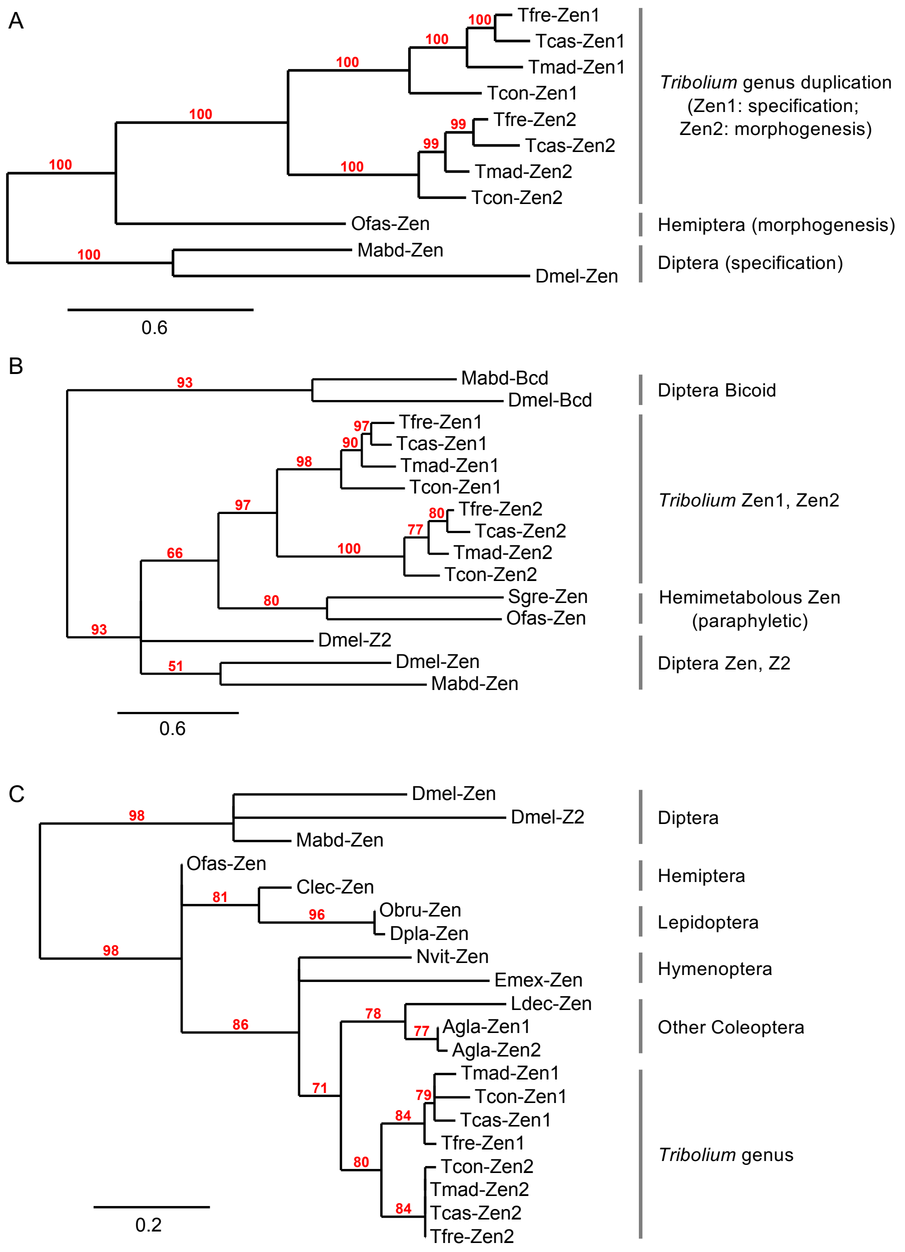


**Figure S2. Additional phylogenies of insect class 3 Hox proteins (Zen and Bicoid orthologues and paralogues).** Phylogenetic trees that differ in taxa, gene class, protein sequence input, and phylogenetic method consistently support an origin of the *zen* tandem duplication within the *Tribolium* genus of the Coleoptera. Taxon and protein sampling choice is for Zen proteins with known extraembryonic functions **(A)**, insect class 3 Hox genes with known expression patterns **(B)**, and wider sampling for taxonomic diversity based on sequence data alone (**C**, two species from each of five insect orders). All *Tribolium* species are included in all analyses. Trees were generated by: Bayesian methods on full-length Zen and Bicoid proteins (A-B), or maximum likelihood with the homeodomain alone (C; compare with full-length proteins and maximum likelihood in Fig. 1B); see main text methods for further details. In total, 38 trees with different parameters and taxa (30 proteins from 21 arthropod species) were analyzed, with consistent results.

Results: Across the trees shown here and Fig. 1B, *Tribolium* Zen proteins form a well-supported clade (≥76%), with subclades for each of Zen1 and Zen2 (≥84%). The topology within each paralogue's subclade also parallels the species tree (Fig. 1A), except where there is a loss of resolution in (C), which does not examine full-length proteins. However, rapid sequence divergence and long branch attraction consistently result in dipteran sequences falling as molecular outgroups, and with deviations between the gene trees and species tree relationships at the ordinal level and for more ancient divergence times.

Abbreviations and sequence sources (GenBank accessions unless otherwise noted): Agla, *Anoplophora glabripennis* independent Zen duplicates (OGS v1.2: Zen1, AGLA004144-RA; Zen2, AGLA014266-RA; [1]); Clec, *Cimex lectularius* (OGS v1.2: CLEC001685-RA; [2]); Dmel, *Drosophila melanogaster* Zen (NP_476793.1), Z2 (NP_476794.1), Bicoid (NP_731111.1); Dpla, *Danaus plexippus plexippus* (OWR47614.1); Emex, *Eufriesea mexicana* (XP_017753571); Ldec, *Leptinotarsa decemlineata* (OGS v1.2: LDEC004404-PA; [3]); Mabd, *Megaselia abdita* Zen (CAB40893.1), Bicoid (CAB40892.1); Nvit, *Nasonia vitripennis* (XP_001603758.2); Obru, *Operophtera brumata* (KOB75729.1); Ofas, *Oncopeltus fasciatus* (ABC17998.1, and OGS v1.2: OFAS000840-RA); Sgre, *Schistocerca gregaria* (CAB61208.1); *Tribolium* species as in the main text. Where protein IDs are from a specified OGS version, these gene models are available at https://i5k.nal.usda.gov/, last accessed 27 February 2020.

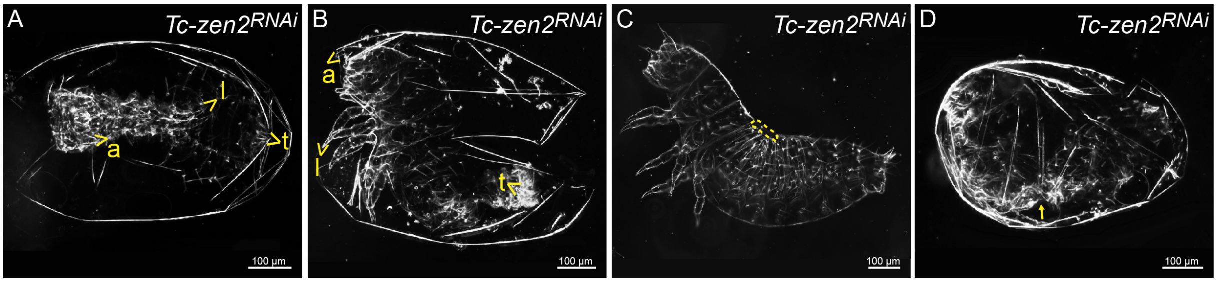


**Figure S3. Phenotypic consequences of *Tc-zen2* RNAi – milder defects, based on larval cuticle preparations. (A)** Partially everted phenotype - anterior inside out. **(B)** Partially everted phenotype - posterior inside out. **(C)** Dorsal open (small medial hole within dashed circle). **(D)** Cuticle crumbs - larval cuticle is constricted in the middle (arrow), presumably due transient contractile squeezing by the EEMs after a loss in tissue integrity [4]. Anatomical abbreviations: a, antenna; l, limb; t, telson.

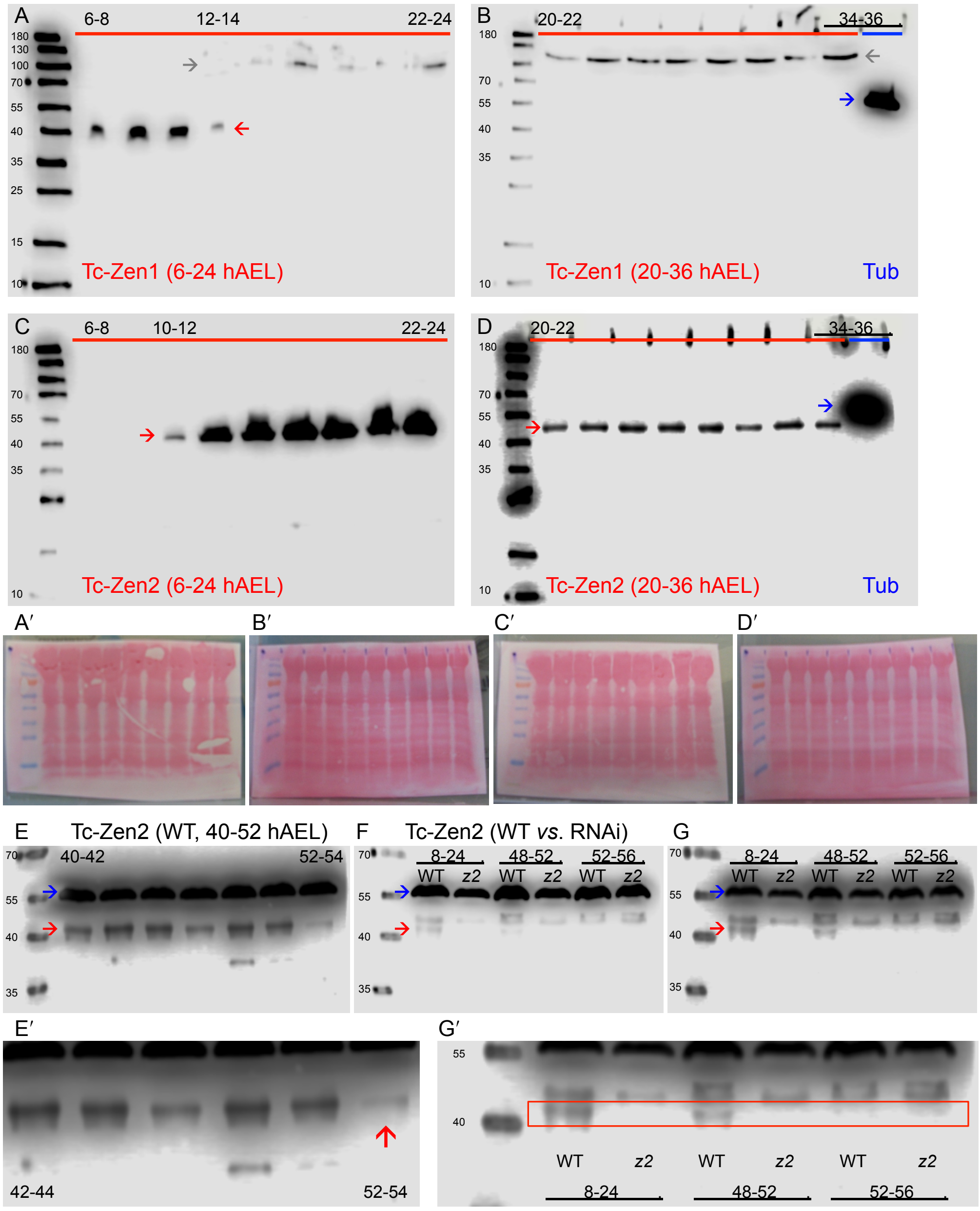


**Figure S4. Tc-Zen1 and Tc-Zen2 expression detected by western blotting.** Images presented here show additional blots in support of Fig. 4A. **(A-D)** Early through mid development expression of Tc-Zen1 (A-B) and Tc-Zen2 (C-D), sampled in two-hour intervals from blastoderm through extended germband stages (A, C: 6-24 hAEL), and extended germband through mid germband retraction stages (B, D: 20-36 hAEL, note overlap of staging between left and right blots). In these samples, the membrane was cut vertically and the strongly expressed anti-Tubulin control (59.0 kDa, blue arrows, "Tub") is only detected in the rightmost lane of the mid-embryogenesis blots (B, D). Both Tc-Zen1 and Tc-Zen2 are cleanly detected as single bands (red arrows). The background signal >100 kDa (grey arrows) in the Tc-Zen1 blots (A-B) is attributed to debris associated with the development of the serosal cuticle (from 14 hAEL) in these whole-egg preparations. **(A′-D′)** Ponceau S staining to visualize whole-egg lysate material. **(E-E′)** Late development expression of Tc-Zen2 (40-54 hAEL). Although Tc-Zen2 expression drops from 50-52 hAEL (pre-rupture), its expression is still detected during membrane withdrawal (arrow at 52-54 hAEL in the enlarged inset image in E′), but it can only be observed on high intensity images (same blot as in Fig. 4A, here shown at higher intensity). **(F-G′)** To confirm RNAi knockdown for *Tc-zen2*, we compared wild type (WT) and RNAi samples (*z2*) at selected stages: across a broad interval of early development when wild type Tc-Zen2 is strongly expressed (8-24 hAEL), and at each of the stages chosen for the late RNA-seq analysis (pre-rupture, 48-52 hAEL; during withdrawal, 52-56 hAEL). The same blot is shown at low (F) and high (G) intensity. Red and blue arrows indicate Tc-Zen2 and anti-Tubulin, respectively. Note the absence of expression of Tc-Zen2 in the RNAi samples, highlighted within the red-boxed region in the high magnification inset in G′. However, as wild type Tc-Zen2 is very weakly expressed from 52 hAEL (E), both wild type and RNAi samples show an absence of Tc-Zen2 at the 52-56 hAEL stage. Nonetheless, as the 52-56 hAEL samples were obtained within the same experiments as the 48-52 hAEL samples, we considered these samples to represent valid RNAi knockdown material, a fact borne out in the subsequent RNA-seq analyses. The bands immediately above the Tc-Zen2 band do not vary by stage or treatment (relative to the anti-Tubulin control) and are therefore interpreted as unspecific background. Left-hand size standards are in kDa (all bands labeled in A, selected bands labeled in subsequent blots).

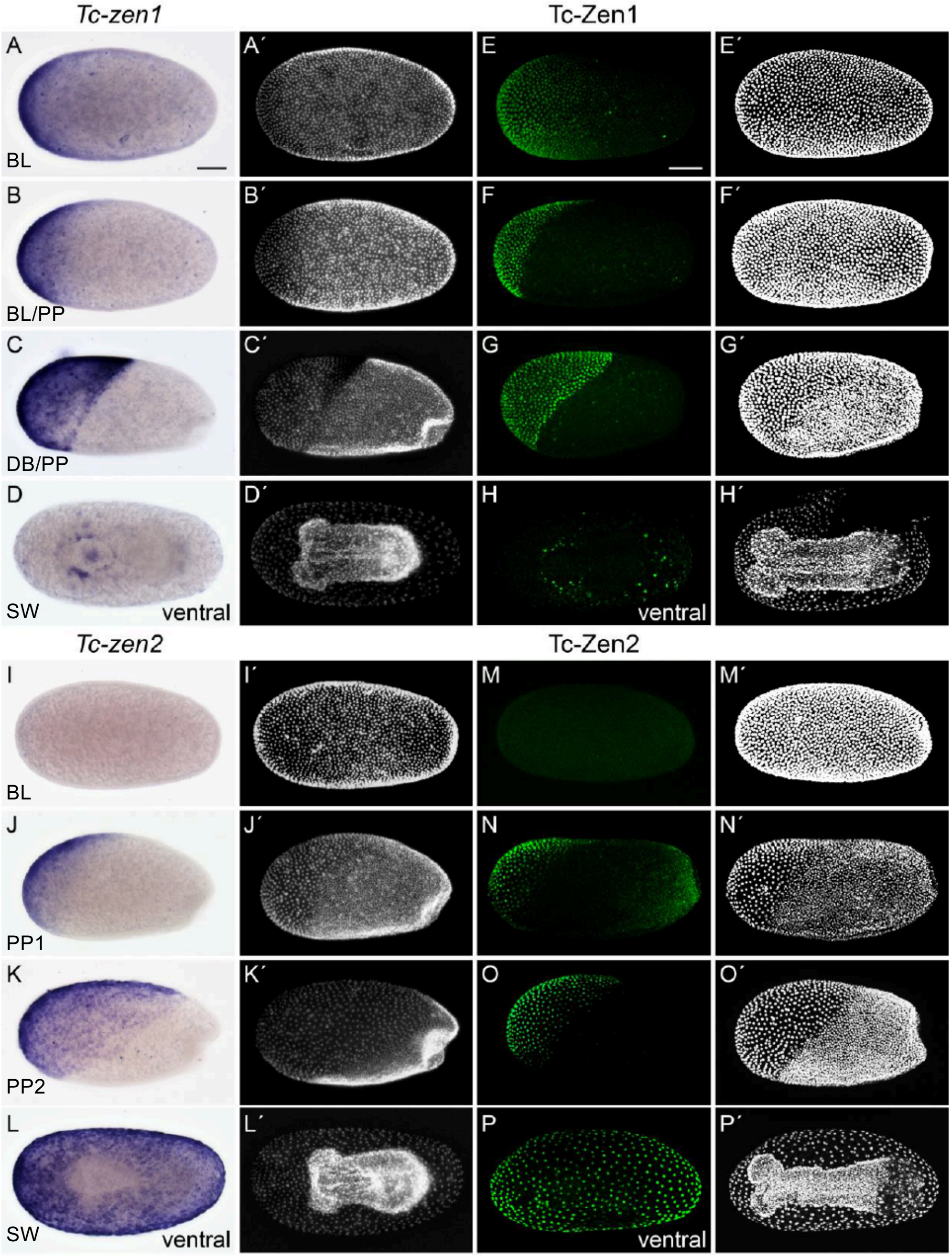


**Fig. S5. Early *in vivo* expression dynamics of transcript and protein for the *Tc-zen* paralogues.** **(A-D)** *Tc-zen1 in situ* hybridization, **(E-H)** Tc-Zen1 immunohistochemistry, **(I-L)** *Tc-zen2 in situ* hybridization, **(M-P)** Tc-Zen2 immunohistochemistry. Whole mount eggs are shown from the blastoderm through serosal window closure stages, in support of Figs. 3-4, using the same *in situ* probes and paralogue-specific peptide antibodies [6] (panels A, I, J, P here are reproductions of main text figure panels 3B, 3G, 3H, and 4C, respectively). Letter-prime images show a nuclear counterstain. Images are oriented with anterior left, in lateral aspect with dorsal up unless otherwise indicated or for uniform blastoderm stages. Scale bars are 100 µm (A, E) and apply to all respective *in situ* and immunohistochemistry images and their counterstains. Staging abbreviations: BL, blastoderm formation/ uniform blastoderm; DB, differentiated blastoderm; PP, primitive pit; SW, serosal window.

Overall, there is a temporal offset whereby *Tc-zen1* precedes *Tc-zen2*. For both genes, protein expression (and nuclear localization) immediately follows transcript expression. By the serosal window stage, *Tc-zen1* has largely switched off (D, H), while *Tc-zen2* has expanded throughout the serosa (L, P).

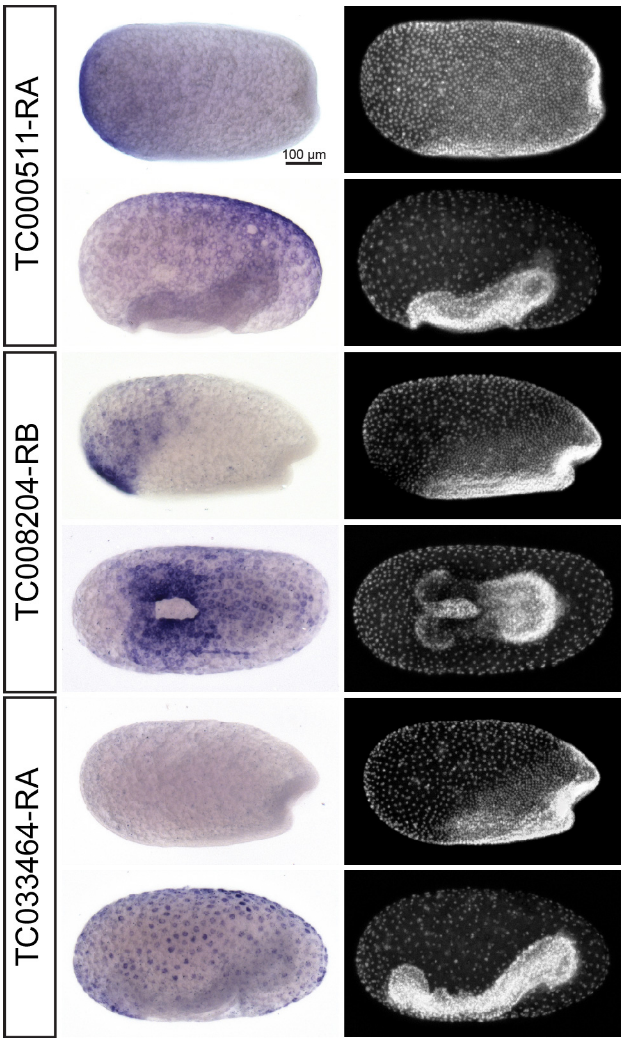


**Figure S6. Expression screen summary of early Tc-Zen2 candidate targets.** Genes were chosen based on RNA-seq after RNAi differential expression ($P_{adj}$≤0.01, |log2 FC|≥1, at 10-14 hAEL; Table S1B), with three of 19 tested genes shown here. For each gene, expression was evaluated by *in situ* hybridization during early and late EEM formation: "primitive pit" (upper image) and "serosal window" (lower image) stages, respectively, with staging determined by DAPI counterstain (right images). Early developmental dynamics mean that these three validated candidates, all with specific serosal expression, include factors that are either upregulated or downregulated by Tc-Zen2, and with distinct staging for the onset of expression. All views are lateral except for a ventral view in the lower panel for *TC008204-RB*; scale bar is 100 µm (shown in first micrograph and applies to all images).

***TC000511-RA*** (log2 FC -1.8074; *Drosophila* homologue *hadley*, peak expression during pupal stages: see comment in Fig. S6) expression starts in the anterior of the serosa. During serosa expansion the expression spreads into the whole serosa, although transcript levels then appear to decline in the most anterior part.

***TC008204-RB*** (log2 FC 1.2187; *Arylalkylamine N-acetyltransferase 1, aaNAT*, cuticle sclerotization factor) expression begins with highest transcript levels detected in the anterior-ventral region of the serosa. This gene is activated by Tc-Zen1 before being repressed by Tc-Zen2 (see Fig. 6I), and this is reflected in the contracting expression domain centered around the serosal rim at the later stage, similar to but broader than the domain of *Tc-zen1* at this stage (*cf.*, Figs. 3D-F,6E).

***TC033464-RA*** (log2 FC -2.7879; *Drosophila* CG13618 homologue, kinase domain) transcript is not detected until the serosal window stage, when perinuclear expression is observed throughout the tissue, consistent with this being a later Tc-Zen2 target during serosal maturation.

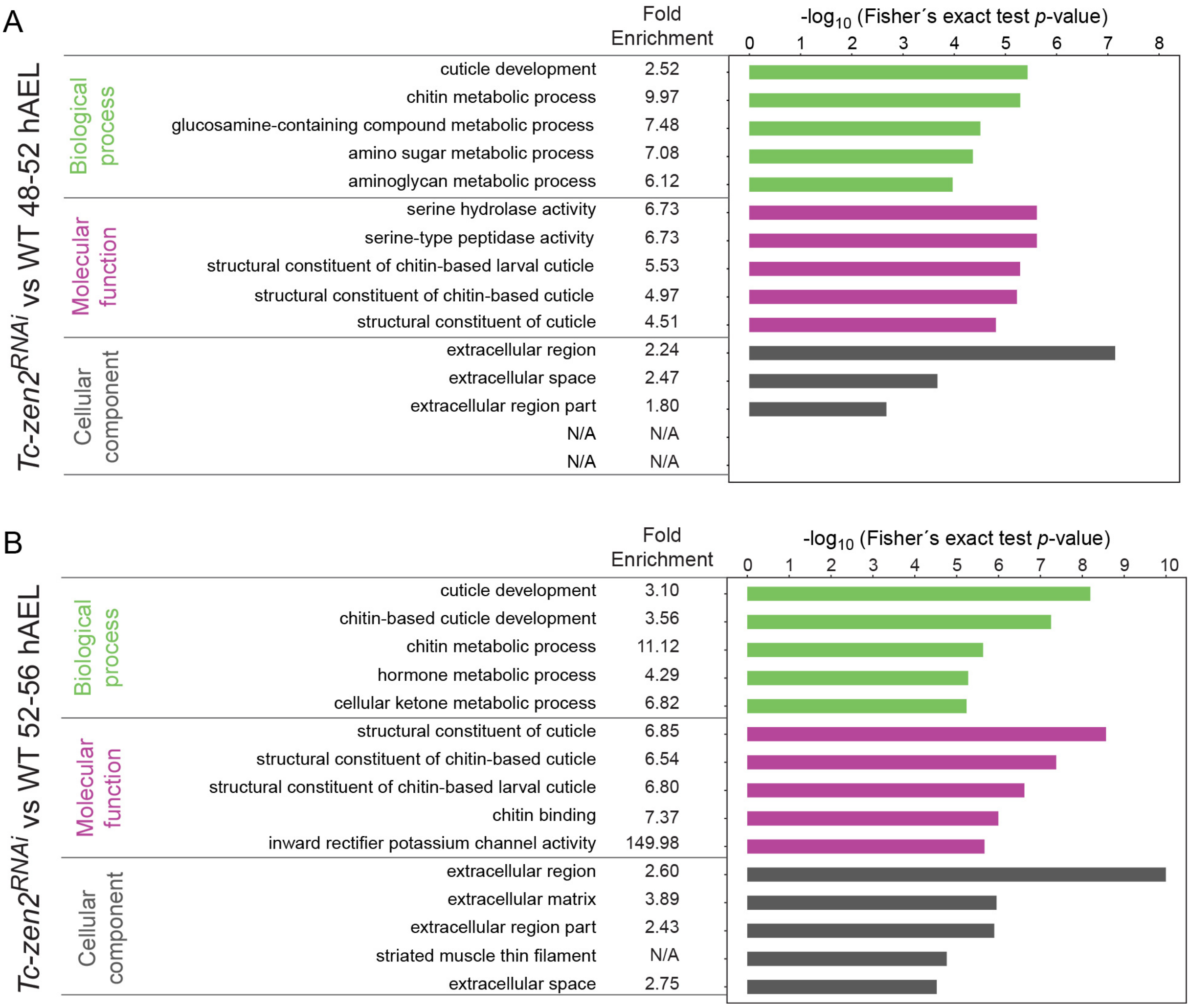


**Figure S7. Gene ontology (GO) enrichment analysis** of the differentially expressed genes in the pre-rupture **(A)** and rupture/post-rupture **(B)** stages after *Tc-zen2* RNAi. If applicable, the five most significantly enriched GO terms and their fold enrichment per GO domain are shown for each of the datasets (Fisher´s exact test, two-sided). Only over-represented GO terms are shown.

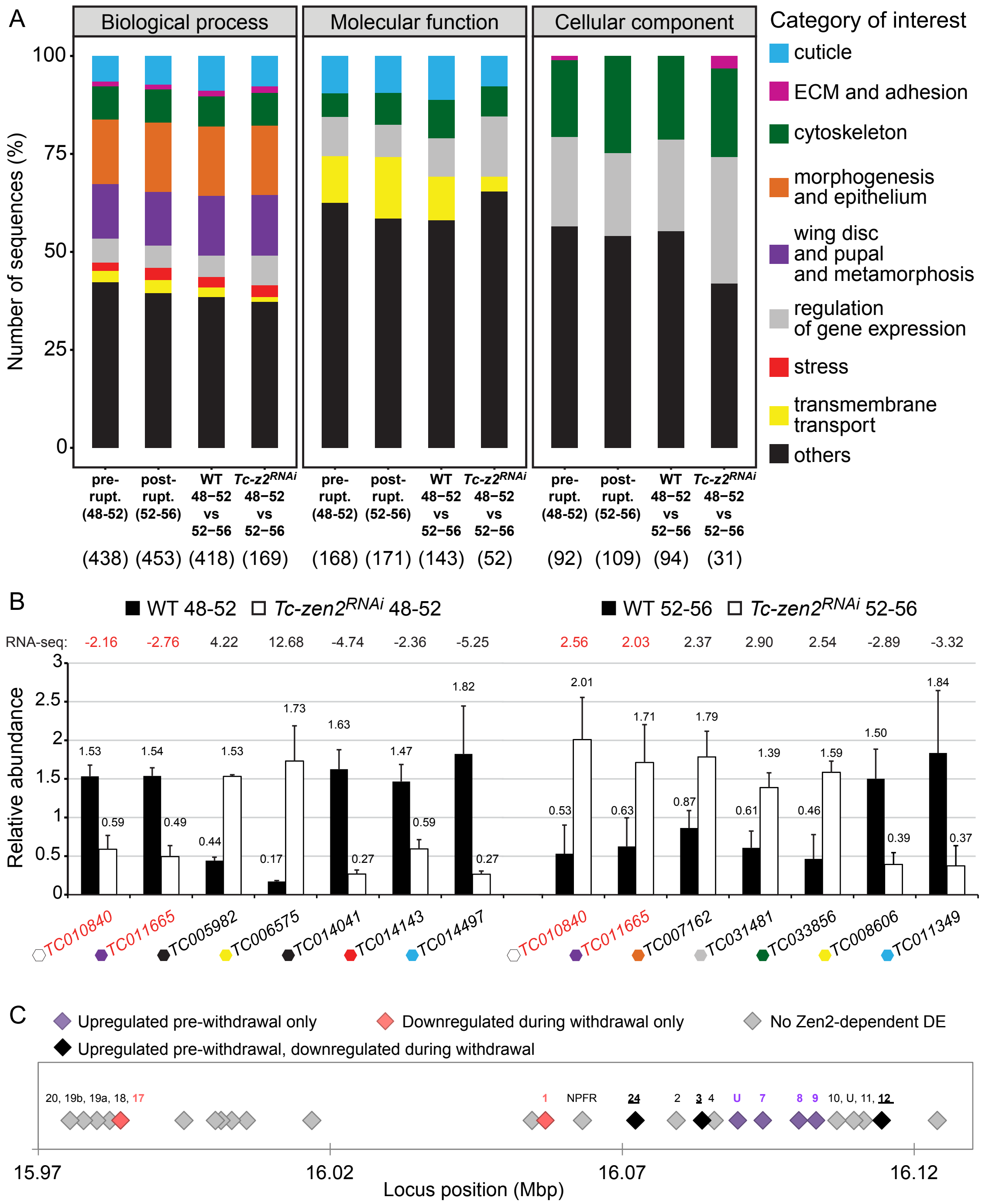


**Figure S8. Functional annotation and validation of late developmental targets of Tc-Zen2.**
**(A)** GO profiles per pairwise DE comparison (as in Fig. 5D). Our GO categories reflect known serosal features (note legend; see main text Methods, Table S5). In addition to cytoskeletal, epithelial, and extracellular remodeling, we considered *Drosophila* imaginal disc morphogenesis, which has noted similarities to EEM withdrawal [4]. We included transcriptional regulation, as *Tc-zen2* may act upstream in serosal gene regulatory networks. We also looked for stress response genes, given the unnatural constraint of the embryo in the absence of EEM withdrawal [7]. These categories account for ~60%, ~35%, and ~50% of sequences for the respective categories biological process, molecular function, and cellular component. Transmembrane transport proteins comprise an additional, prominent category that we had not deliberately selected (yellow). Transporters may serve physiological roles in the serosa as a barrier epithelium, with the potential to mediate exchange between the egg and the outer environment and for yolk catabolism [8-10]. Numbers are GO domains

per DE comparison. Note that a given gene may be assigned to multiple GO categories, depending on available information in public databases. **(B)** RT-qPCR testing of selected Tc-Zen2 candidate targets. Candidates were chosen from the GO categories in A (note color-coding, where white indicates a novel gene; see also Table S6) and based on the direction of Tc-Zen2 regulation. Seven candidates were chosen from each stage (48-52 hAEL and 52-56 hAEL), where two of these show changing direction of regulation across stages (red text). All RNA-seq DE predictions (fold change values above the graph) were validated by RT-qPCR (mean value from three biological replicates; error bars represent one standard deviation). **(C)** Mapping of Tc-Zen2-dependent regulation to genes within the *Osiris* complex of tandemly duplicated, co-expressed *Osiris* genes (shown for a portion of scaffold NC_007424.3, based on Official Gene Set v3 (OGS3), gene locus positions as defined in http://ibeetle-base.uni-goettingen.de, last accessed 5 August 2019). All predicted genes are represented by diamonds, with the *Osiris* gene homology indicated above *Osiris* family genes (U: uncertain homology) and for the conserved, syntenic factor NPFR [11], while unrelated genes are unlabeled. Notably, although the *Osiris* genes are co-expressed and thus represent an exemplar locus of open, accessible chromatin for co-regulation, we detect Tc-Zen2-dependent regulation only for a subset of this gene family. Moreover, whereas some tandem genes are co-regulated by Tc-Zen2 (purple: *Osiris genes U/7/8/9*), other genes that are co-regulated and show changing Tc-Zen2-dependent regulation occur throughout the gene complex (black: *Osiris 24, 3, 12*). As detailed in Table S6, gene TC011665, validated by RT-qPCR in panel B, is *Osiris 12*. Regulation color-coding is as in main text Figure 5E. Staging is the same as in the rest of the figure (48-52 hAEL: pre-withdrawal; 52-56 hAEL: during withdrawal morphogenesis).

**Table S5. List of gene ontology (GO) terms grouped by category of interest in each GO domain.**

| Category of interest | GO domain: Biological process |
|---|---|
| Stress | response to endoplasmic reticulum stress |
| | cellular response to oxidative stress |
| | regulation of translation in response to stress |
| | regulation of response to osmotic stress |
| | stress-activated protein kinase signaling cascade |
| | positive regulation of stress fiber assembly |
| | age-dependent response to oxidative stress |
| | regulation of stress fiber assembly |
| Cuticle | chitin-based cuticle development |
| | molting cycle, chitin-based cuticle |
| | cuticle pigmentation |
| | chitin-based cuticle sclerotization |
| | ecdysis, chitin-based cuticle |
| | regulation of adult chitin-containing cuticle pigmentation |
| | regulation of chitin-based cuticle tanning |
| | cuticle pattern formation |
| Cytoskeleton | cytoskeleton organization |
| | actin cytoskeleton organization |
| | actin filament organization |
| | actin filament bundle assembly |
| | cytoskeleton-dependent cytokinesis |
| | actin filament-based movement |
| | regulation of actin cytoskeleton organization |
| | establishment or maintenance of cytoskeleton polarity |
| | cytoskeleton-dependent intracellular transport |
| | microtubule cytoskeleton organization involved in mitosis |
| | cytoskeletal anchoring at plasma membrane |
| | oocyte microtubule cytoskeleton organization |
| | establishment or maintenance of microtubule cytoskeleton polarity |
| | regulation of actin filament length |
| Epithelium and morphogenesis | epithelium development |
| | morphogenesis of an epithelium |
| | epithelial tube morphogenesis |
| | cell morphogenesis |
| | cell part morphogenesis |
| | cell projection morphogenesis |
| | imaginal disc morphogenesis |
| | post-embryonic animal organ morphogenesis |
| | imaginal disc-derived appendage morphogenesis |
| | post-embryonic appendage morphogenesis |
| | cell morphogenesis involved in differentiation |
| | sensory organ morphogenesis |
| | epithelial cell development |
| | gland morphogenesis |
| | digestive tract morphogenesis |
| | epithelium migration |
| | regulation of organ morphogenesis |
| | Malpighian tubule morphogenesis |
| | dendrite morphogenesis |
| | morphogenesis of embryonic epithelium |
| | embryonic hindgut morphogenesis |
| | regulation of cell morphogenesis |
| | regulation of morphogenesis of an epithelium |
| | antennal morphogenesis |
| | trachea morphogenesis |
| | epithelial cell proliferation involved in renal tubule morphogenesis |
| | epithelial tube formation |
| | mesoderm morphogenesis |
| | transepithelial transport |
| | spermathecum morphogenesis |
| | heart morphogenesis |

| | |
|---|---|
| Epithelium and morphogenesis | branching morphogenesis of an epithelial tube |
| | negative regulation of cell morphogenesis involved in differentiation |
| | morphogenesis of a branching epithelium |
| | chaeta morphogenesis |
| | regulation of cell proliferation involved in imaginal disc-derived wing morphogenesis |
| | male genitalia morphogenesis |
| | imaginal disc-derived leg joint morphogenesis |
| | intestinal epithelial cell differentiation |
| | positive regulation of cell morphogenesis involved in differentiation |
| | post-embryonic genitalia morphogenesis |
| | cell elongation involved in imaginal disc-derived wing morphogenesis |
| Wing disc and pupal and metamorphosis | instar larval or pupal development |
| | wing disc development |
| | instar larval or pupal morphogenesis |
| | metamorphosis |
| ECM and adhesion | cell-matrix adhesion |
| | extracellular matrix organization |
| | negative regulation of cell-cell adhesion |
| Regulation of gene expression | gene expression |
| | regulation of gene expression |
| | negative regulation of gene expression |
| | positive regulation of gene expression |
| | regulation of translational initiation |
| | chromatin silencing |
| | negative regulation of chromatin silencing |
| | regulation of chromatin silencing |
| Transmembrane transport | transmembrane transport |
| | regulation of transmembrane transport |
| | regulation of transmembrane transporter activity |
| | positive regulation of ion transmembrane transporter activity |
| | positive regulation of ion transmembrane transport |
| **Category of interest** | **GO domain: Molecular function** |
| Regulation of gene expression | DNA binding |
| | transcription factor activity, RNA polymerase II distal enhancer sequence-specific binding |
| | transcription factor activity, RNA polymerase II core promoter proximal region sequence-specific binding |
| | transcriptional activator activity, RNA polymerase II transcription regulatory region sequence-specific binding |
| | transcriptional repressor activity, RNA polymerase II transcription regulatory region sequence-specific binding |
| | flavin adenine dinucleotide binding |
| | purine ribonucleotide binding |
| | purine nucleoside binding |
| | purine nucleotide binding |
| | ribonucleoside binding |
| | purine ribonucleoside triphosphate binding |
| | RNA binding |
| | repressing transcription factor binding |
| | regulatory region nucleic acid binding |
| | chromatin insulator sequence binding |
| Cuticle | structural constituent of chitin-based larval cuticle |
| | chitin deacetylase activity |
| Cytoskeleton | myosin binding |
| | actin binding |
| | actinin binding |
| | actin filament binding |
| Transmembrane transport | ion transmembrane transporter activity |
| | secondary active transmembrane transporter activity |
| | primary active transmembrane transporter activity |
| | organic acid transmembrane transporter activity |
| | sulfate transmembrane transporter activity |
| | active ion transmembrane transporter activity |

| | |
|---|---|
| Transmembrane transport | monoamine transmembrane transporter activity |
| | taurine transmembrane transporter activity |
| | phosphate ion transmembrane transporter activity |
| | serotonin transmembrane transporter activity |
| **Category of interest** | **GO domain: Cellular component** |
| Regulation of gene expression | nucleus |
| | chromatin |
| | nuclear chromosome part |
| | nuclear chromosome |
| | chromosomal region |
| | nuclear transcription factor complex |
| Cytoskeleton | cytoskeleton |
| | contractile fiber |
| | unconventional myosin complex |
| | polymeric cytoskeletal fiber |
| | cell cortex region |
| | cell cortex part |
| | myosin II complex |
| ECM and adhesion | focal adhesion |

**Table S6. Description of Tc-Zen2 late candidate target genes validated by RT-qPCR.** *Dmel* - *Drosophila melanogaster*; FC - fold change. See Fig. S6B.

| gene ID | developmental stage | GO descriptor | *Dmel* homologue | FC |
|---|---|---|---|---|
| ***TC010840*** | pre-rupture<br>post-rupture | no information available;<br>DUF4773 (pfam 15998) | no homologue | -2.16<br>2.56 |
| ***TC011665*** | pre-rupture<br>post-rupture | morphogenesis and epithelia (imaginal disc-derived wing morphogenesis);<br>DUF1676 (pfam 07898) | Osiris 12 | -2.76<br>2.03 |
| ***TC005982*** | pre-rupture | cytosol (other) | Major facilitator superfamily transporter 3 | 4.22 |
| ***TC006575*** | pre-rupture | transmembrane transport | Pickpocket 26 | 12.68 |
| ***TC014041*** | pre-rupture | cell differentiation (other) | Z band alternatively spliced PDZ-motif | -4.74 |
| ***TC014143*** | pre-rupture | stress | Protein kinase, cAMP-dependent, catalytic subunit 3 | -2.36 |
| ***TC014497*** | pre-rupture | cuticle | Cuticular protein 65Av | -5.25 |
| ***TC007162*** | post-rupture | morphogenesis and epithelia | Ribbon | 2.37 |
| ***TC031481*** | post-rupture | regulation of gene expression | Nuclear factor interleukin-3-regulated protein-like protein | 2.90 |
| ***TC033856*** | post-rupture | cytoskeleton | CG8213 | 2.54 |
| ***TC008606*** | post-rupture | transmembrane transport | Excitatory amino acid transporter 3-like protein | -2.89 |
| ***TC011349*** | post-rupture | cuticle | Chondroitin proteoglycan 2-like protein | -3.32 |

**Table S7.** ***Tribolium castaneum*** **(*TC*) primer sequences for *in situ* hybridization, RNAi, and RT-qPCR.** Lower case letters in the primer sequences indicate adapter sequences for subsequent amplification with the T7 promoter universal primers, as described [12].

| TC gene identifier / primer orientation | sequence | amplicon size cDNA/gDNA [bp] |
|---|---|---|
| ***in situ* hybridization** | | |
| *TC000921* (*Tc-zen1*) / F | ggccgcggTCCCAATTTGAAAACCAAGC | 688 |
| *TC000921* (*Tc-zen1*) / R | cccggggcCGTTCCACCCTTCCTGATAA | |
| *TC000922* (*Tc-zen2*) / F | ggccgcggAACGCCCCAGTTTTCAACAA | 546 |
| *TC000922* (*Tc-zen2*) / R | cccggggcCTCATCCTTCACCACCACCT | |
| **RNAi** | | |
| *TC000921* (*Tc-zen1*) / F | ggccgcggTTTGAAAACCAAGCCGTTCT | 203 (short fragment) |
| *TC000921* (*Tc-zen1*) / R | cccggggcCGTTGGGGTTGAGTTTCTTG | |
| *TC000921* (*Tc-zen1*) / F | ggccgcggTTTGAAAACCAAGCCGTTCT | 682 (long fragment) |
| *TC000921* (*Tc-zen1*) / R | cccggggcCGTTCCACCCTTCCTGATAA | |
| *TC000922* (*Tc-zen2*) / F | ggccgcggCAATGTCGCCGCAATCGACG | 250 |
| *TC000922* (*Tc-zen2*) / R | cccggggcACACAATTCTTCCCTTGGTA | |
| **RT-qPCR** | | |
| *TC000921* (*Tc-zen1*) / F | TCCACCTTCTGATTGGAACTG | 105/161 |
| *TC000921* (*Tc-zen1*) / R | CGTTGGGGTTGAGTTTCTTG | |
| *TC000922* (*Tc-zen2*) / F | TCGAAGTGTCCCTCTCAGAAA | 101/147 |
| *TC000922* (*Tc-zen2*) / R | GGAGGAGGTGTACGCAGTTC | |
| *TC008261* (*Tc-RpS3*) / F | ACCGTCGTATTCGTGAATTGAC | 186 |
| *TC008261* (*Tc-RpS3*) / R | ACCTCAAAACACCATAGCAAGC | |
| *TC010840-RA* / F | TGCGCCTCTCTTCAGTACCT | 127/563 |
| *TC010840-RA* / R | CGCCAGGTAAAGGCATACAC | |
| *TC011665-RA* / F | AAGACGCAGCTTTGACCAAT | 142/198 |
| *TC011665-RA* / R | ATCATCATACCGCCCATCAT | |
| *TC005982-RA* / F | GTATTGTGTACGCGGGGACT | 117/2790 |
| *TC005982-RA* / R | TTGTTGAAAACCCACCCTCT | |
| *TC006575-RA* / F | TTACCATTTGTCCCGAGTCC | 144/193 |
| *TC006575-RA* / R | CGAACTTCGTGTCGCAAATA | |
| *TC014041-RA* / F | TATGGCAGCCACAAGAAGC | 150/5369 |
| *TC014041-RA* / R | GTTGGGGTGGTGTCGTAGAT | |
| *TC014143-RA* / F | CGAAGACGATAAAGAGGGCTA | 123/5213 |
| *TC014143-RA* / R | TTCATGGCACTATACTGGTTCG | |
| *TC014497-RA* / F | GCTCTTCGTTTCACTTGTGG | 137/183 |
| *TC014497-RA* / R | TGCCGTCACTGGTCTCATAC | |
| *TC007162-RA* / F | CGTCGCAACCTGTAAGTCTG | 148/3480 |
| *TC007162-RA* / R | TCGTTCATCAGCGTGAAGTC | |
| *TC008606-RA* / F | CAAGCTGGCCTCGTCACTAT | 134/10182 |
| *TC008606-RA* / R | ATGCAGTCGCACATCACATT | |
| *TC011349-RA* / F | ACCATCGTTACCCTCATTGC | 122423 |
| *TC011349-RA* / R | TCCCTGCATTCGATATAGCC | |
| *TC031481-RA* / F | GCACCCCGATAACGGATTA | 114/4865 |
| *TC031481-RA* / R | GGGATTTCACCATTTACTGGA | |
| *TC033856-RA* / F | GAAGAGGCCGAAAACTACGA | 150/1157 |
| *TC033856-RA* / R | GCCCCTTTACCACCGACTAT | |

**Table S8. Comparison of alignment statistics by read length**. Data are from all the sequenced samples collected during early development. The difference in mapping efficiency of the same sample mapped as 75-bp reads and 100-bp reads is shown. The samples mapped as 75-bp reads have higher mapping efficiency, gaining 9-15% of uniquely mapping reads and only 3-4% of multi-mapping reads. Overall alignment rate represents the sum of uniquely mapping and multi-mapping reads.

| | Unique-mapping [%] | | Multi-mapping [%] | | Overall alignment rate [%] | |
|---|---|---|---|---|---|---|
| **Sample name** | **100 bp** | **75 bp** | **100 bp** | **75 bp** | **100 bp** | **75 bp** |
| *Tc-zen1*$^{RNAi}$ 1 | 55 | 68 | 14 | 17 | 69 | 85 |
| *Tc-zen1*$^{RNAi}$ 2 | 55 | 69 | 13 | 17 | 68 | 86 |
| *Tc-zen1*$^{RNAi}$ 3 | 54 | 64 | 15 | 19 | 69 | 83 |
| *Tc-zen1* WT 1 | 51 | 66 | 14 | 18 | 65 | 84 |
| *Tc-zen1* WT 2 | 54 | 68 | 14 | 17 | 68 | 85 |
| *Tc-zen1* WT 3 | 56 | 67 | 14 | 17 | 70 | 84 |
| *Tc-zen2*$^{RNAi}$ 1 | 57 | 68 | 14 | 17 | 71 | 85 |
| *Tc-zen2*$^{RNAi}$ 1 | 58 | 68 | 14 | 17 | 72 | 85 |
| *Tc-zen2*$^{RNAi}$ 1 | 57 | 68 | 14 | 17 | 71 | 85 |
| *Tc-zen2* WT 1 | 56 | 64 | 15 | 18 | 71 | 82 |
| *Tc-zen2* WT 2 | 58 | 67 | 14 | 17 | 72 | 84 |
| *Tc-zen2* WT 3 | 55 | 66 | 14 | 17 | 69 | 83 |
| **range** | **51-58** | **64-69** | **13-15** | **17-19** | **65-72** | **82-86** |
| **difference** | **9-15** | | **3-4** | | **11-19** | |

#### References


1. McKenna DD, Scully ED, Pauchet Y, Hoover K, Kirsch R, Geib SM, Mitchell RF, Waterhouse RM, Ahn S-J, Arsala D, et al: Genome of the Asian longhorned beetle (*Anoplophora glabripennis*), a globally significant invasive species, reveals key functional and evolutionary innovations at the beetle–plant interface. *Genome Biol* 2016, 17:227.
2. Benoit JB, Adelman ZN, Reinhardt K, Dolan A, Poelchau M, Jennings EC, Szuter EM, Hagan RW, Gujar H, Shukla J, et al: Unique features of a global human ectoparasite identified through sequencing of the bed bug genome. *Nat Commun* 2016, 7:10165.
3. Schoville SD, Chen YH, Andersson MN, Benoit JB, Bhandari A, Bowsher JH, Brevik K, Cappelle K, Chen MM, Childers AK, et al: A model species for agricultural pest genomics: the genome of the Colorado potato beetle, *Leptinotarsa decemlineata* (Coleoptera: Chrysomelidae). *Sci Rep* 2018, 8:1931.
4. Hilbrant M, Horn T, Koelzer S, Panfilio KA: The beetle amnion and serosa functionally interact as apposed epithelia. *eLife* 2016, 5:e13834.
5. Gasteiger E, Hoogland C, Gattiker A, Duvaud S, Wilkins MR, Appel RD, Bairoch A: Protein identification and analysis tools on the ExPASy server. In *The Proteomics Protocols Handbook.* Edited by Walker JM. Totowa, New Jersey: Humana Press; 2005: 571-607
6. Mackrodt D: Etablierung und Funktion maternaler Proteingradienten im *Tribolium* Blastoderm. *Ph.D. (Dr. rer. nat.).* Friedrich-Alexander-Universität Erlangen-Nürnberg, Naturwissenschaftlichen Fakultät; 2016.
7. Panfilio KA: Late extraembryonic development and its *zen-RNAi*-induced failure in the milkweed bug *Oncopeltus fasciatus*. *Dev Biol* 2009, 333:297-311.
8. Dorn A: Ultrastructure of embryonic envelopes and integument of *Oncopeltus fasciatus* Dallas (Insecta, Heteroptera) I. Chorion, amnion, serosa, integument. *Zoomorphologie* 1976, 85:111-131.
9. Lamer A, Dorn A: The serosa of *Manduca sexta* (Insecta, Lepidoptera): ontogeny, secretory activity, structural changes, and functional considerations. *Tissue Cell* 2001, 33:580-595.
10. Panfilio KA: Extraembryonic development in insects and the acrobatics of blastokinesis. *Dev Biol* 2008, 313:471-491.
11. Shah N, Dorer DR, Moriyama EN, Christensen AC: Evolution of a large, conserved, and syntenic gene family in insects. *G3 (Bethesda)* 2012, 2:313-319.
12. Koelzer S, Kölsch Y, Panfilio KA: Visualizing late insect embryogenesis: Extraembryonic and mesodermal enhancer trap expression in the beetle *Tribolium castaneum*. *PLoS One* 2014, 9:e103967.
